# svaseq: removing batch effects and other unwanted noise from sequencing data

**DOI:** 10.1101/006585

**Authors:** Jeffrey T. Leek

## Abstract

It is now well known that unwanted noise and unmodeled artifacts such as batch effects can dramatically reduce the accuracy of statistical inference in genomic experiments. We introduced surrogate variable analysis for estimating these artifacts by (1) identifying the part of the genomic data only affected by artifacts and (2) estimating the artifacts with principal components or singular vectors of the subset of the data matrix. The resulting estimates of artifacts can be used in subsequent analyses as adjustment factors. Here I describe an update to the sva approach that can be applied to analyze count data or FPKMs from sequencing experiments. I also describe the addition of supervised sva (ssva) for using control probes to identify the part of the genomic data only affected by artifacts. These updates are available through the surrogate variable analysis (sva) Bioconductor package.

## 1 Background

Batch effects and other technological artifacts introduce spurious correlation, create bias, and add variability to the results of genomic experiments [1, 2, 3]. Unfortunately all the possible sources of bias are unknown in most high-throughput experiments [4, 5]. In some cases, it is possible to rely on the date the samples were processed as a surrogate for unmeasured artifacts [6]. However, each new technology may suffer from different artifacts and it may take time for the community to discover which variables must be measured and included in an analysis [7].

In 2007 we introduced surrogate variable analysis as a conceptual approach to statistical modeling of genomic data when artifacts are unknown or unmeasured [4] and subsequently improved the estimation algorithm [8] (Figure 1). The conceptual idea we proposed modeled the data as a combination of known variables of interest, known adjustment variables, and unknown and unmeasured artifacts. A simple version of this model might relate gene expression for gene *i* on sample *j* (*g*_*ij*_) to the phenotype for that sample (*y*_*j*_), the known batch variable for that sample (*a*_*j*_) and an unknown artifact on that sample (*u*_*j*_):

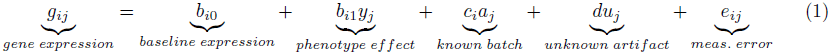

**Figure 1:**
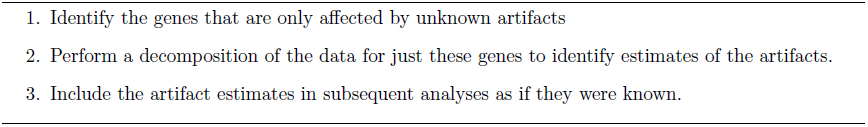
Surrogate variable analysis. The general surrogate variable analysis framework for identifying unknown artifacts in genomic data has three steps [4, 8].

**Figure 2:**
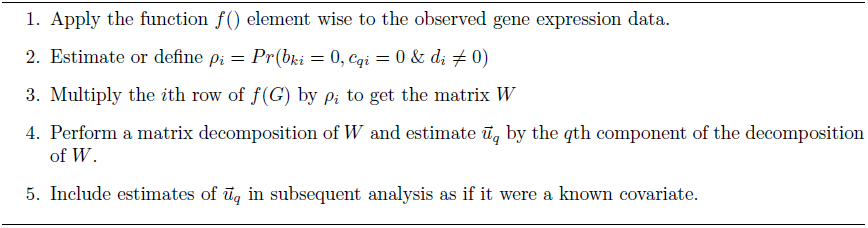
General surrogate variable analysis estimation framework. In this general framework, Step 1 allows for transformations specific to different data types, Step 2 allows for either estimating or defining the probabilities of being affected by unknown artifacts but not known variables, and Step 4 allows for a variety of matrix decompositions and factor analysis approaches.

**Figure 3:**
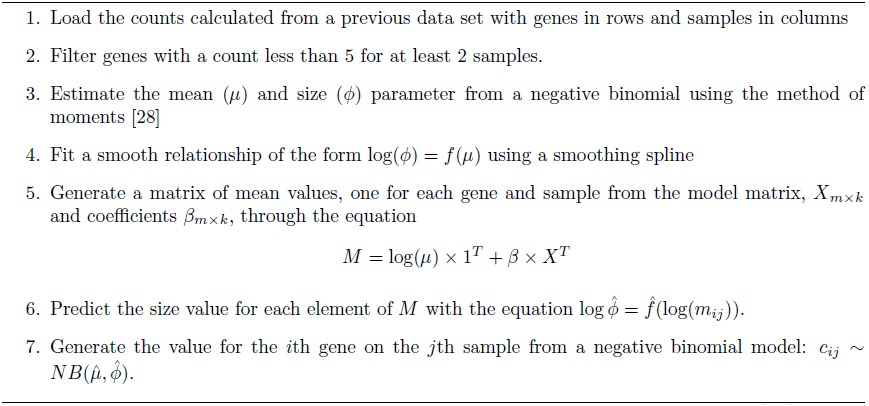
Approach for simulating RNA-seq data with Polyester package [29].

If only a single gene is measured, it is difficult to estimate the unknown artifact *u*_*j*_ from the data directly, since all the coefficients (*b*, *c*, *d*) are unknown. But we noticed [4] that if many genes are measured, it is possible that for some genes the coefficients for *b*_1_*_i_* and *c*_*i*_ may be equal to zero. For these specific genes the model reduces to:

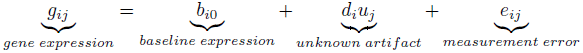

Our next insight was that even though *d*_*i*_ and *u*_*j*_ are unknown, you do not need to know either of them exactly to get correct statistical inference for the parameters for the phenotype variable *b*_*i*1_, you just have to know their linear combination *d*_*i*_ × *u*_*j*_ [4, 8]. We showed that if you collect the data for all genes where there is no effect of phenotype or known batch (*b*_1_*_i_* = 0 and *c*_*i*_ = 0) and subtracted the mean of each gene to remove the baseline effect, the matrix form of the model for the data is:

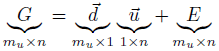

where *m*_*u*_ is the number of genes where *b*_1_*_i_* = 0 and *c*_*i*_ = 0 and *n* is the sample size. We then pointed out that you could apply a matrix decomposition like the singular value decomposition or principal components analysis to this subset of the data to estimate surrogates for *G* and *u*. Later we also showed that if the number of genes that were not affected by phenotype or known batch (*b*_1_*_i_* = 0 and *c* = 0) but were affected by unknown artifacts (*d*_*i*_ ≠ 0) was large enough, you can obtain consistent estimates of a linear transformation of *g*_*i*_ [9]. The surrogate variable analysis approach then proceeds in three steps:

In this paper I discuss two new extensions of the surrogate variable analysis approach for sequencing data inspired by the work of other groups. The first extension, which I call supervised surrogate variable analysis (ssva) was introduced by Gagnon-Bartsch and colleagues [5]. The idea is to perform Step of the surrogate variable analysis algorithm (Figure 1). The second idea, svaseq, uses a moderated log link in place of the identity link when estimating the surrogate variables in Steps 1-2. This is a simplified version of a similar extension of surrogate variable analysis first proposed by Risso and colleagues [10].

## 2 Methods

### 2.1 General form of the surrogate variable analysis mathematical model

The general form of the simple model in equation (1) is:

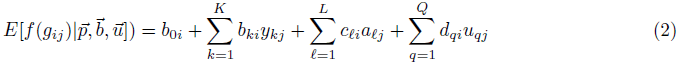

or in matrix form:

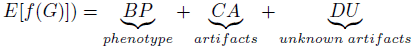

where the function *f* () has been applied component wise to each element of *G* and there may be multiple phenotypes, artifacts, or unknown artifacts. The general surrogate variable analysis algorithm then proceeds in the following four concrete steps.

### 2.2 Relationship of surrogate variable analysis to other approaches

The general form of the surrogate variable analysis algorithm relies on the idea that there is a subset of genes, probes, or transcripts that are affected by unknown heterogeneity but are not affected by the biological relationship of interest. This is the main way that a surrogate variable analysis approach is distinguished from a standard principal component regression. Standard principal component based regression methods, such as EIGENSTRAT [11], are sufficient when the number of probes, genes, or features expected to show a signal is small. Then the principal components will entirely capture artifacts and not the signal of interest.

A number of extensions and variations on the surrogate variable analysis algorithm have been introduced. For step one in the sva algorithm, identifying probes only associated with batch, it has been proposed to use control probes [5, 10]. For step two of the sva algorithm, estimating latent factors only associated with dependence, it has been proposed to use factor analysis [12], independent components analysis [13], and principal components analysis [14]. Another extension of the surrogate variable approach in step two has been to model known sources of technical or biological covariation between the measurements for probes, for example in eQTL studies [15, 16].

### 2.3 Supervised sva (ssva)

ySupervised sva (ssva) sets *p*_*i*_ = 1 for all negative controls and *p*_*i*_ = 0 for all other genes in Step 2 of the sva algorithm. The assumption is that control probes will capture all of the variation due to unknown artifacts and none of the variation due to the phenotype. Control probes may miss biological artifacts. For example, we showed that trans-eQTL that are associated with multiple gene expression levels may act like an artifact when measuring the association between gene expression and phenotype [4]. These artifacts may be missed by the ssva approach. However ssva is particularly useful for unfortunate experimental designs where the phenotype variable and unknown artifacts are highly correlated [6], making empirical estimates unstable [5].

### 2.4 Moderated log link sva (svaseq)

The second extension involves the choice of function *f* () in (1). In our original work we used the identify function for data measured on an approximately symmetric and continuous scale. For sequencing data, which are often represented as counts, a more suitable model may involve the use of a moderated log function [17, 18]. For example in Step 1 of the algorithm we may first transform the gene expression measurements by applying the function *log*(*g*_*ij*_ + *c*) for a small positive constant. In the analyses that follow I will set *c* = 1. After performing Steps 1-5 of the surrogate variable analysis estimation algorithm, the estimated covariates are included in downstream models as adjustment variables. For the analyses that follow, I will use the limma package [19] with the voom method [20] for differential expression analysis.

### 2.5 Combining svaseq and ssva

Supervised svaseq proceeds by applying the transformation *log*(*g*_*ij*_ + *c*) to the gene expression count data in Step 1 and setting *p*_*i*_ = 1 for all negative controls and *p*_*i*_ = 0 for all other genes in Step 2 of the sva algorithm.

### 2.6 Zebrafish data

I use data from Zebrafish sensory neurons with three control samples and three gallein treated samples as the comparison groups [21]. These data are available as part of the *zebrafishRNAseq* Bioconductor package. I loaded the data and filtered as described in the RUVSeq package. Then I estimated batch effects using supervised and unsupervised surrogate variable analysis for sequencing, principal components analysis, RUV with control probes, RUV with empirical controls, and residual RUV. I compared the model estimates and I also compared differential expression analysis results when each of the different batch effect estimates was included in the model in place of the study variable.

### 2.7 ReCount data

ReCount is a database of pre-processed RNA-sequencing data, processed to be comparable across samples [22]. In this analysis, I downloaded pre-counted RNA-sequencing data sets measuring gene expression in two separate Hapmap populations [23, 24]. For our analysis, I downloaded the count data from ReCount and downloaded the pedigree information from the Hapmap website. I then performed differential expression analyses looking for differences in expression between males and females and estimated unknown latent structure. I calculated estimates of batch effects using unsupervised surrogate variable analysis for sequencing, principal components analysis, RUV with empirical control probes, and RUV on residuals. I compared the estimates to the variable indicating whether the data came from the Pickrell or Montgomery study. I compared two scenarios, one where the sex and study variables were balanced and one where they were imbalanced. I also compared the differential expression analysis results when each of the different batch effect estimates was included in the model in place of the study variable.

### 2.8 GEUVADIS data

I downloaded the processed GEUVADIS [25, 26] Ballgown object [27] from https://github.com/alyssafrazee/ballgown_code

I also compared the differential expression analysis results when each of the different batch effect estimates was included in the model in place of the laboratory variable.

### 2.9 Simulating Data

I simulated data from a negative binomial model for count based RNA-sequencing data. I used the following algorithm to simulate data, for complete details see the simulated data R markdown document and accompanying HTML file.

I estimated the model parameters from the Zebrafish data described above. I simulated two scenarios, one where the group and batch variable where balanced and one where they were imbalanced. Data were simulated with the *Polyester* R package [29]. Then I estimated batch effects using supervised and unsupervised surrogate variable analysis for sequencing, principal components analysis, RUV with control probes, RUV with empirical controls, and residual RUV. I compared the model estimates to the true simulated batch variable and I also compared differential expression analysis results when each of the different batch effect estimates was included in the model in place of the study variable.

### 2.10 Code and availability

ssva and svaseq are currently implemented in the sva Bioconductor package version 3.11.2 or greater (http://www.bioconductor.org/packages/devel/bioc/html/sva.html). All data and code used to perform this analysis are available as R markdown files [30] available from: https://github.com/jtleek/svaseq. You can view the individual analyses as webpages at:

1. Zebrafish analysis: http://jtleek.com/svaseq/zebrafish.html
2. ReCount analysis: http://jtleek.com/svaseq/recount.html
3. GEUVADIS analysis: http://jtleek.com/svaseq/geuvadis.html
4. Simulated data analysis: http://jtleek.com/svaseq/simulateData.html

## 3 Results

### 3.1 Simulated data

I estimated simulation parameters from the zebrafish data as described in the methods. I then performed several checks to confirm that the data generated by the simulated model recapitulated the qualitative behavior of the data used to estimate the model parameters (Figure (4)), that data generated without signal did not show statistically significant results, data could be simulated with differential expression signal, and that data with batch effects displayed the expected conservative bias of p-values [31] (See supplementary analysis files http://jtleek.com/svaseq/zebrafish.html).

**Figure 4:**
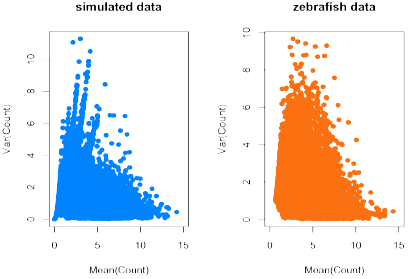
Distribution of means and variances for simulated and real zebrafish data. To confirm that my simulation procedure produced reasonable simulated counts, I plotted the gene-specific means and variances for (a) the simulated data set and (b) the observed zebrafish data set. The two distributions are qualitatively similar. Additional checks on the simulation procedure are provided in the simulated data analysis at http://jtleek.com/svaseq/simulateData.html.

In this case there was a single simulated batch effect. I simulated two scenarios, in the first scenario the batch effect and the group effect had low correlation (). As expected, all methods that aim to estimate batch effects while taking into account multiple sources of signal (svaseq and RUV methods) are highly correlated with the simulated batch effect. The estimate of batch based on principal components is biased, because the principal component is estimating a linear combination of the group and batch variable (Figure 5).

**Figure 5:**
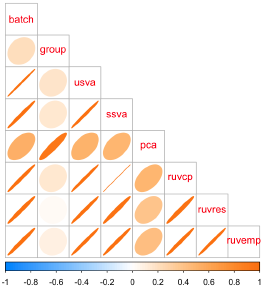
Correlation between simulated batch and group variables and various batch estimates. Light circles indicate low correlation and dark, tight ellipses indicate high correlation. In this case, all estimates that respect multiple sources of signal (sva and RUV based) methods are highly correlated with the simulated batch effect. Principal components estimates a linear combination of the group and batch variable and has lower concordance with the true simulated batch and the other estimates. Additional details at http://jtleek.com/svaseq/simulateData.html.

**Figure 6:**
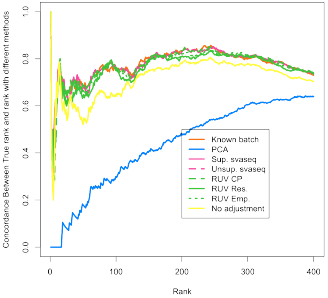
Differential expression results for simulated data. A concordance at the top plot (CAT plot) shows the fraction of DE results that are concordant between the analysis with the true batch and the analyses using different batch estimates. Supervised (pink solid) and unsupervised (pink dotted) surrogate variable analysis for sequencing, RUV with control probes (green dashed), RUV with empirical controls (green dotted), and residual RUV (green solid) all outperform not adjusting for batch effects (yellow) while principal components analysis (blue) performs worse than no adjustment. Additional details at http://jtleek.com/svaseq/simulateData.html.

**Figure 7:**
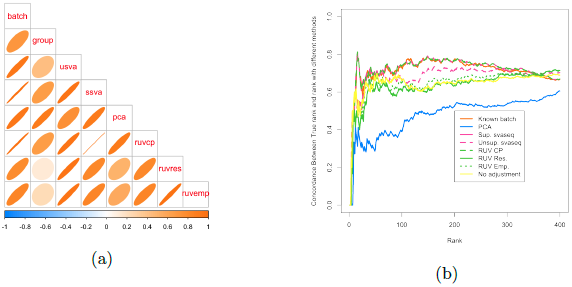
Comparison of batch effect results when group and batch are correlated. (a) A plot of the correlation between the different batch estimates and the batch variable analogous to Figure 5. (b) A concordance at the top plot measuring concordance between the analysis using the true batch variable and the various estimates analogous to Figure 6. Here the unsupervised RUV approaches using empirical control probes and residuals perform worse than no adjustment, because the methods can not distinguish signal from the known group variable and the unknown batch variable. Additional details at http://jtleek.com/svaseq/simulateData.html.

**Figure 8:**
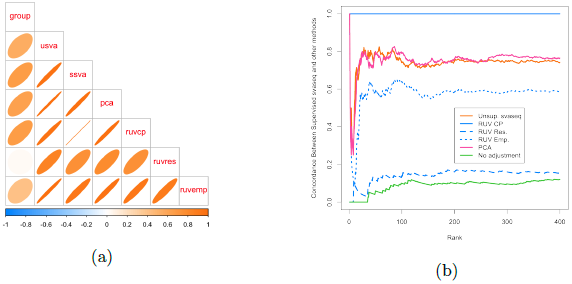
Comparison of batch effect results on zebrafish data. (a) A plot of the correlation between the different batch estimates analogous to Figure 5, but with no gold standard. (b) A concordance at the top plot measuring concordance between the analysis using the supervised SVA estimates and the various other batch estimates analogous to Figure 6. The control probes RUV approach (blue solid in (b)) and supervised sva approach produce identical results. The unsupervised sva (orange solid) and principal components (pink solid) approaches are most similar to the supervised estimates in this scenario. Additional details at http://jtleek.com/svaseq/zebrafish.html.

I next fit models relating the gene expression counts to the simulated group phenotype (*p*) and batch effect estimates (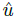) using the following model:

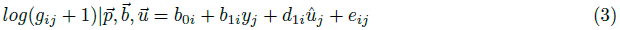

I accounted for the potential relationship between mean and variance using the voom method [20]. I then estimated how concordant the differential expression results were with the results we obtained when we fit model 6 using the true simulated batch variable using concordance at the top plots (CAT plots) [32]. For each ranking, these plots show the fraction of results that are the same between the analysis using the true batch variable and the batch variable estimated with different methods. Supervised and unsupervised surrogate variable analysis for sequencing, RUV with control probes, RUV with empirical controls, and residual RUV all outperform not adjusting for batch effects while principal components analysis performs worse than no adjustment. The reason is that the principal component estimate is correlated with group and absorbs some of the signal due to that variable.

We performed an identical analysis where the batch was now correlated with the group variable. Qualitatively similar results hold in this second simulated scenario with one exception. The empirical RUV methods attempt to define control probes by identifying genes that do not show differential expression with respect to batch. But when batch and group are correlated, this may also through away genes that show signal with respect to the group variable. Similar the residual RUV approach estimates the batch variable after taking the residuals from the model fit of the counts on the group variable. However, when batch and group are correlated, this again removes batch signal and leads to slightly lower performance of the RUV approaches [4]. Unsupervised svaseq does not use the control probes but avoids some of these difficulties by iteratively identifying probes associated with group but not associated with batch [8] (Figure 9).

**Figure 9:**
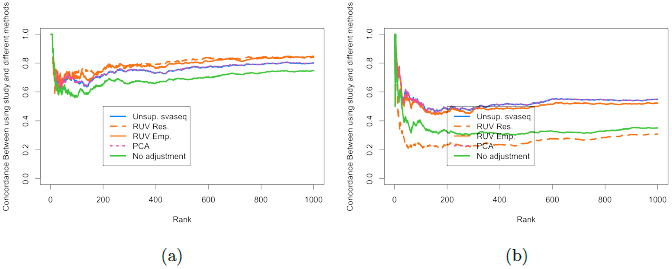
Comparison of differential expression results for ReCount experiment. (a) A concordance at the top plot measuring concordance between the analysis using the true study and the various other batch estimates analogous to Figure 6. (b) A concordance at the top plot measuring concordance between the analysis using the true study and the various other batch estimates analogous to Figure 6 when data were resampled to make the sex and study variables moderately correlated (*r*^2^ = 0.33.) When sex and study are uncorrelated, RUV performs slightly better and when sex and study are correlated, svaseq performs slightly better. Additional details at http://jtleek.com/svaseq/recount.html

### 3.2 Zebrafish data

Next I performed an analysis on the zebrafish data as described in the methods section. Here, the batch variable is not known, but we do have negative control probes which can be used to estimate the batch effects. When comparing the batch estimates, I noted that the supervised sva estimates and the RUV control probes estimates were perfectly correlated (correlation 1) and that they produced identical differential expression results. The unsupervised sva and principal components approaches are most similar to the supervised estimates from SVA or RUV for the zebrafish data.

### 3.3 ReCount data

For the ReCount data I generated an artificial batch effect by combining the data from two different studies of gene expression in two different populations [23, 24]. I used sex as the outcome variable in the analysis and then estimated batch effects using the same set of proposed approaches. I next fit models relating the gene expression counts (*g*) to sex variable (*p*, phenotype variable representing sex) and batch effect estimates (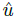, representing the study the samples came from) using the following model:

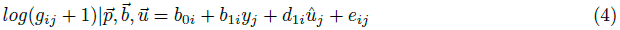

I accounted for the potential relationship between mean and variance using the voom method [20]. I then estimated how concordant the differential expression results were with the results we obtained when we fit model 6 using the true simulated batch variable using concordance at the top plots (CAT plots) [32].

In the original data, the batch effect and the group variable are nearly perfectly orthogonal. In this situation the empirical and residual RUV approaches produce estimates of the batch variable more highly correlated than the unsupervised svaseq approach (see http://jtleek.com/svaseq/recount.html) and produce correspondingly more similar differential expression results to using the true study variable as an adjustment in the differential expression analysis (Figure **??**a). However, I next re-sampled the data to mimic a scenario where the group and batch variable showed modest correlation (*r*^2^ = 0.33). In this scenario the unsupervised sva and principal components analysis approaches outperform the empirical control RUV approach. The residual RUV approach performs worse than no adjustment for study, because signal due to the batch variable was removed when the residuals from the model relating sex to phenotype was calculated (Figure **??**b).

### 3.4 GEUVADIS data

The *Ballgown* R package https://github.com/alyssafrazee/ballgown [27] can be used to analyze abundance data from assembled transcriptome data from *Cufflinks* [33]. I loaded data from the GEUVADIS project [25, 26] that we recently processed using *Cufflinks* and *Ballgown* [27]. I selected only the non-duplicated samples and performed a differential expression analysis comparing different populations. I then compared the estimated batch effects using the various approaches to the known lab where the samples were processed, one of the variables that showed the highest association with assembled transcript levels [26].

I assessed concordance between the batch effect estimates and the lab variable by fitting the model:

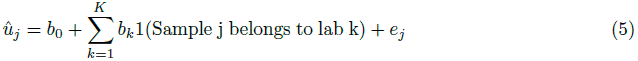

and then performed an ANOVA to compare the model including lab to the null model of no association with lab. The unsupervised sva and principal components estimates showed significantly higher F-statistics for concordance (482 and 456, respectively) compared to the RUV approach (106 and 109 for RUV residual and empirical, respectively). I next fit models relating the gene expression counts (*g*) to the population phenotype (*p*, phenotype variable representing population) and batch effect estimates (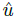, representing the study the samples came from) using the following model:

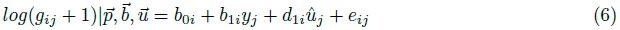

I compared the results to the differential expression model where I included the known lab variable as an adjustment in place of 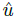. The svaseq and principal components adjusted analyses showed greater concordance with the lab adjusted analysis, as expected since the batch estimates were more highly correlated with this known variable (Figure 10).

**Figure 10:**
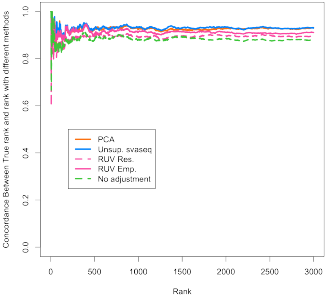
Differential expression results for GEUVADIS data. A concordance at the top plot (CAT plot) shows the fraction of DE results that are concordant between the analysis with the true laboratory and the analyses using different batch estimates. Unsupervised surrogate variable analysis for sequencing (blue) and principal components analysis (orange) outperform the RUV based methods (pink) and no batch adjustment (green). Additional details at http://jtleek.com/svaseq/geuvadis.html.

## 4 Discussion

Here I have described the general surrogate variable analysis framework and I have introduced two extensions of the surrogate variable analysis approach. The first takes advantage of known control probes to simplify the surrogate variable analysis algorithm and the second addresses the distribution of count and FPKM data typically observed in sequencing experiments. I have shown that these approaches perform comparably to other batch effect estimation procedures for sequencing when the group and unknown batch variables are uncorrelated and outperform other approaches when the batch and group variable are correlated. These extensions are currently available from the devel branch of the surrogate variable analysis software http://bioconductor.org/packages/devel/bioc/html/sva.html and all analyses are fully reproducible and available as R markdown documents from https://github.com/jtleek/svaseq.

## 5 Acknowledgements

JL is supported by NIH R01 GM10570502 and GM083084.

